# The *Salmonella* LysR family regulator, RipR, activates the SPI-13 encoded itaconate degradation cluster

**DOI:** 10.1101/648865

**Authors:** Steven J. Hersch, William Wiley Navarre

## Abstract

Itaconate is a dicarboxylic acid that inhibits the isocitrate lyase enzyme of the bacterial glyoxylate shunt. Activated macrophages have been shown to produce itaconate, suggesting that these immune cells may employ this metabolite as a weapon against invading bacteria. Here we demonstrate that, *in vitro*, itaconate can exhibit bactericidal effects under acidic conditions similar to the pH of a macrophage phagosome. In parallel, successful pathogens including *Salmonella* have acquired a genetic operon encoding itaconate degradation proteins, which are induced heavily in macrophages. We characterize the regulation of this operon by the neighbouring gene, *ripR*, in specific response to itaconate. Moreover, we develop an itaconate biosensor based on the operon promoter that can detect itaconate in a semi-quantitative manner and, when combined with the *ripR* gene, is sufficient for itaconate-regulated expression in *E. coli*. Using this biosensor with fluorescence microscopy, we observe bacteria responding to itaconate in the phagosomes of macrophage and provide additional evidence that interferon-γ stimulates macrophage itaconate synthesis and that J774 mouse macrophages produce substantially more itaconate than the human THP-1 monocyte cell line. In summary, we examine the role of itaconate as an antibacterial metabolite in mouse and human macrophage, characterize the regulation of *Salmonella*’s defense against it, and develop it as a convenient itaconate biosensor and inducible promoter system.

**Importance:** In response to invading bacteria, immune cells can produce a molecule called itaconate, which can inhibit microbial metabolism. Here we show that itaconate can also directly kill *Salmonella* when combined with moderate acidity, further supporting itaconate’s role as an antibacterial weapon. We also discover how *Salmonella* recognizes itaconate and activates a defense to degrade it, and we harness this response to make a biosensor that detects the presence of itaconate. This biosensor is versatile, working in *Salmonella enterica* or lab strains of *Escherichia coli*, and can detect itaconate quantitatively in the environment and in immune cells. By understanding how immune cells kill bacteria and how the microbes defend themselves, we can better develop novel antibiotics to inhibit pathogens such as *Salmonella*.

## Introduction

The mammalian immune system includes a multitude of weapons to defend against invading microbes and successful pathogens have evolved a plethora of mechanisms to evade, manipulate, or even benefit from these immune responses. One such pathogen, *Salmonella enterica s*erovar Typhimurium (hereafter referred to as *Salmonella*), has acquired a number of *Salmonella* pathogenicity islands (SPI) that support its survival inside of a host organism. For instance, *Salmonella* employs SPI-1 to invade non-phagocytic cells, and SPI-2 allows the bacteria to survive intracellularly – including in macrophage – which is important for *Salmonella* virulence ^1–4^. These traits allow *Salmonella* to invade the gut epithelium and induce intestinal inflammation resulting in the characteristic gastroenteritis disease. Moreover, the induced inflammation is not merely a threat that *Salmonella* must survive, but it has adapted to thrive in the oxidative environment of the inflamed intestine and utilize inflammation-derived metabolites to outcompete resident microbiota ^5–7^.

Itaconate (2-Methylenesuccinic acid, 2-Methylidenebutanedioic acid) is a metabolite originally recognized in fungal species such as *Aspergillus terreus* and produced commercially for use in polymer production ^8–10^. As an unsaturated diacaboxylate that is somewhat similar in structure to succinate, itaconate is a potent inhibitor of the glyoxylate shunt enzyme AceA (isocitrate lyase) ^11–13^. As such, itaconate inhibits bacterial growth on carbon sources such as acetate and fatty acids, conditions that necessitate the glyoxylate shunt to generate the 4-carbon skeletons that are critical for amino acid biosynthesis and central metabolism. Interestingly, it has been demonstrated that activated macrophage employ the IRG1 protein to produce itaconate, with higher concentrations being produced by mouse macrophage than human ^13–15^. Moreover, IRG1 closely associated with vesicles containing *Legionella pneumophila* and itaconate showed bactericidal activity against this pathogen *in vitro* ^15^. Itaconate was also found to inhibit *Salmonella* growth by reducing media pH and itaconate levels in *Salmonella*-infected mice correlated with splenomegaly ^16^. Cumulatively, these works suggest that itaconate acts as a weaponized metabolite that the immune system employs to inhibit the growth of, or kill, invading bacteria.

If itaconate is an immune-derived antibacterial metabolite then it follows logically that successful pathogens must have methods to evade its effects. Indeed an operon has been identified in *Yersinia* (*ripABC* for ‘required for intracellular proliferation’) that encodes three enzymes catalyzing the ATP/succinyl-CoA-dependent degradation of itaconate into pyruvate and acetyl-CoA ^17^. The operon is not restricted to *Yersinia* and a variety of other bacteria including *Pseudomonas* encode homologs or functional analogs of these enzymes. Several lineages of *Salmonella enterica* harbor a cluster of genes (e.g. genes STM3120-STM3117 in S. Typhimurium strain LT2) within SPI-13 that we refer to here as the ‘itaconate response operon’ (IRO). Interestingly, the IRO genes of *Salmonella* have been shown to be induced heavily in macrophage but not under any other condition tested, supporting a role in degrading macrophage-produced itaconate ^18–20^. High throughput screens have suggested that genes from this operon are important for *Salmonella* survival in mice ^21–23^. Furthermore, it has also been shown that SPI-13 is present in many generalist *S. enterica* serovars but not in some human-restricted serovars of *Salmonella* (e.g. serovars Typhi and Paratyphi A, which encode SPI-8) possibly due to reduced itaconate synthesis by human macrophages ^24^.

In this work we show that itaconate is bactericidal at low but not neutral pH and elucidate the regulation of the *Salmonella* IRO and its induction in mouse and human macrophage. We show that the promoter of the IRO (P_IRO_) is specifically induced by itaconate and that the LysR family transcriptional regulator encoded by the upstream gene, STM3121 (which we propose to name *ripR*), is both necessary and sufficient for this induction. Furthermore, using P_IRO_ with a GFP reporter, we develop a semi-quantitative itaconate biosensor and employ it to show that the IRO is induced heavily in the J774 mouse macrophage cell line but requires interferon-γ (IFN-γ) stimulation to show a detectable response in the THP-1 human monocyte cell line.

## Results

### The *Salmonella* IRO is induced specifically by itaconate in a RipR-dependent manner

The *Salmonella* pathogenicity island-13 includes genes encoding RipC/Ccl (STM3120 in strain LT2), RipB/Ich (STM3119) and RipA/Ict (STM3118), whose homologs have been demonstrated to degrade itaconate into pyruvate and acetyl-CoA ^17^. This operon, which we refer to as the itaconate response operon (IRO), appears to also include the STM3117 gene (encoding a predicted glyoxalase-domain containing protein) and is adjacent to the STM3121 (*ripR*) gene on the reverse DNA strand (Figure 2).

**Figure 1:**
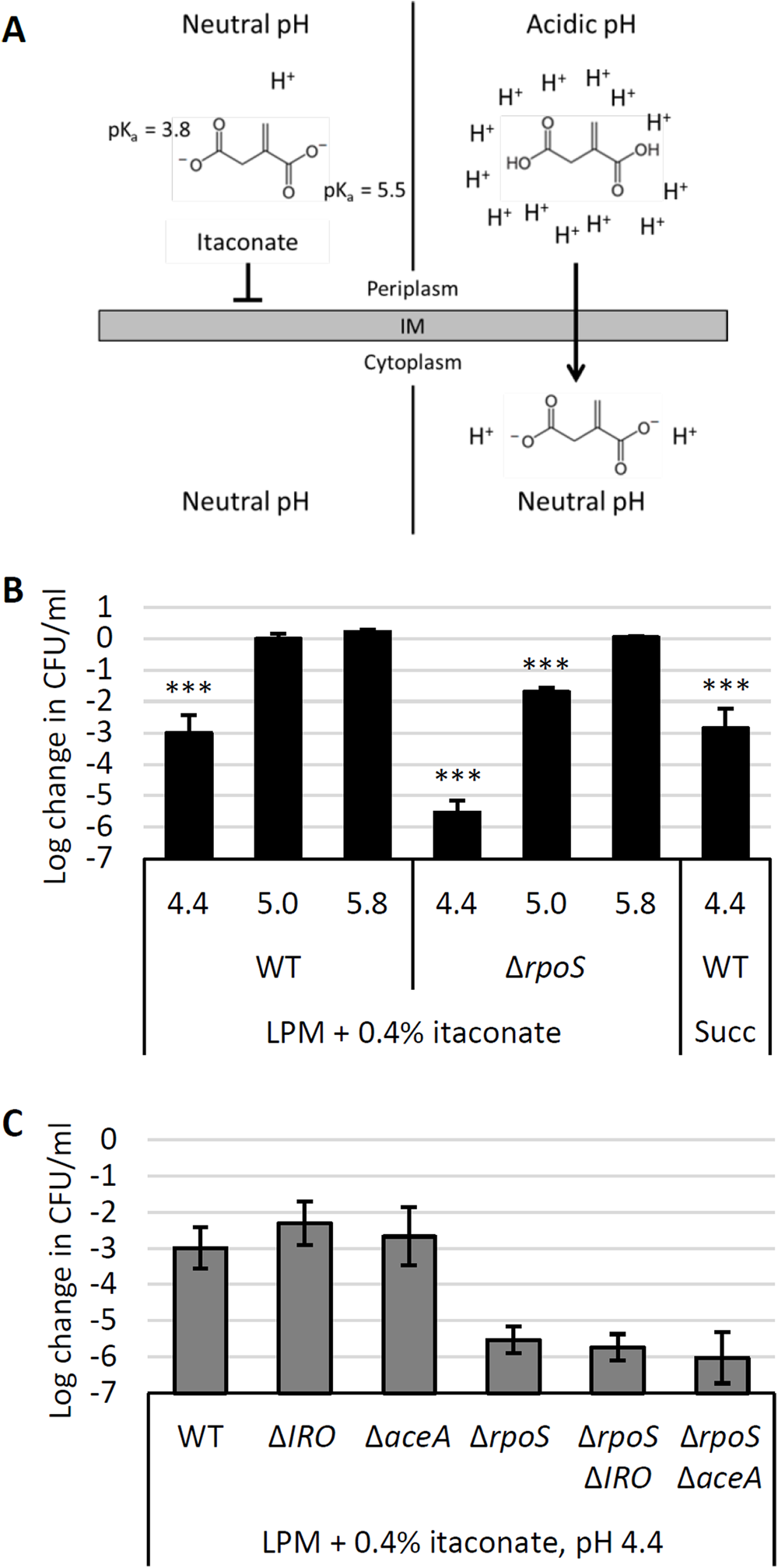
Itaconate is bactericidal at low pH. **A)** At neutral pH, the charges on itaconate inhibit diffusion across a lipid membrane (left). At acidic pH, itaconate protonates to itaconic acid, which can traverse the membrane (right). In the cytoplasm, itaconic acid dissociates and releases protons, trapping it in the cell and acidifying the cytoplasm. **B)** Survival (relative to 0h time point) of wild-type (WT) or *rpoS* mutant *Salmonella* after 3h in LPM media supplemented with 0.4% itaconate or succinate (Succ) and adjusted to pH 4.4, 5.0, or 5.8 as indicated. **C)** As in A but showing survival of *Salmonella aceA* and itaconate response operon (IRO) mutants in LPM + itaconate media at pH 4.4. All data are the average of at least three biological replicates and error bars show one standard deviation. A one-way ANOVA with Sidak’s multiple comparison test was conducted comparing each sample to the same strain at pH 5.8 (B). WT in succinate was compared to WT at pH 5.8 in itaconate. Mutants in panel C were not significantly different than their parental strain (WT or Δ*rpoS*). ***, p < 0.001.

**Figure 2:**
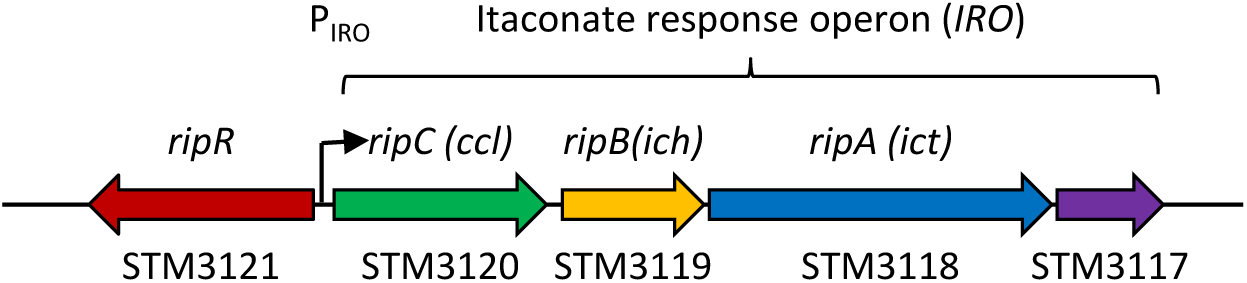
Overview of the itaconate degradation operon (IRO) and the itaconate-responsive regulator, *ripR*. *Salmonella* gene loci based on the genome of *Salmonella* Typhimurium LT2 are shown below. Identities of *ccl, ich*, and *ict* are shown above as previously reported ^17^. STM3117 appears to be encoded in the same operon however its function remains unknown. In this work we designate the names ‘*ripR*’ for STM3121, ‘itaconate response operon’ (IRO) for STM3120-STM3117, and ‘P_IRO_’ for the IRO promoter.

The IRO is strongly induced in cultured mouse macrophages but, according to expression data from SalCom and other published work, is seemingly independent of known virulence regulators (PhoP, RpoS, SsrA/B, OmpR, SlyA, etc.). We hypothesized that itaconate may act as an inducer of IRO expression via the adjacent LysR-family regulator encoded by STM3121. To assess this, we constructed a plasmid-borne fusion of the operon’s promoter (P_IRO_) to superfolder GFP (sfGFP) as a reporter^25^. Indeed, we found that the P_IRO_ promoter was induced highly in the presence of itaconate and this response was entirely dependent on the presence of RipR as neither a *Salmonella ripR* deletion mutant nor the same reporter plasmid in *E. coli* K12 (which does not encode *ripR*) showed induction (Figure 3A). In contrast, when the *ripR* gene was included on the reporter plasmid, itaconate-induced P_IRO_ expression was restored in both the Δ*ripR Salmonella* strain and in *E. coli*. These data not only demonstrate that P_IRO_ is induced in response to itaconate, but also that the neighbouring gene, *ripR*, is both necessary and sufficient for this induction.

**Figure 3:**
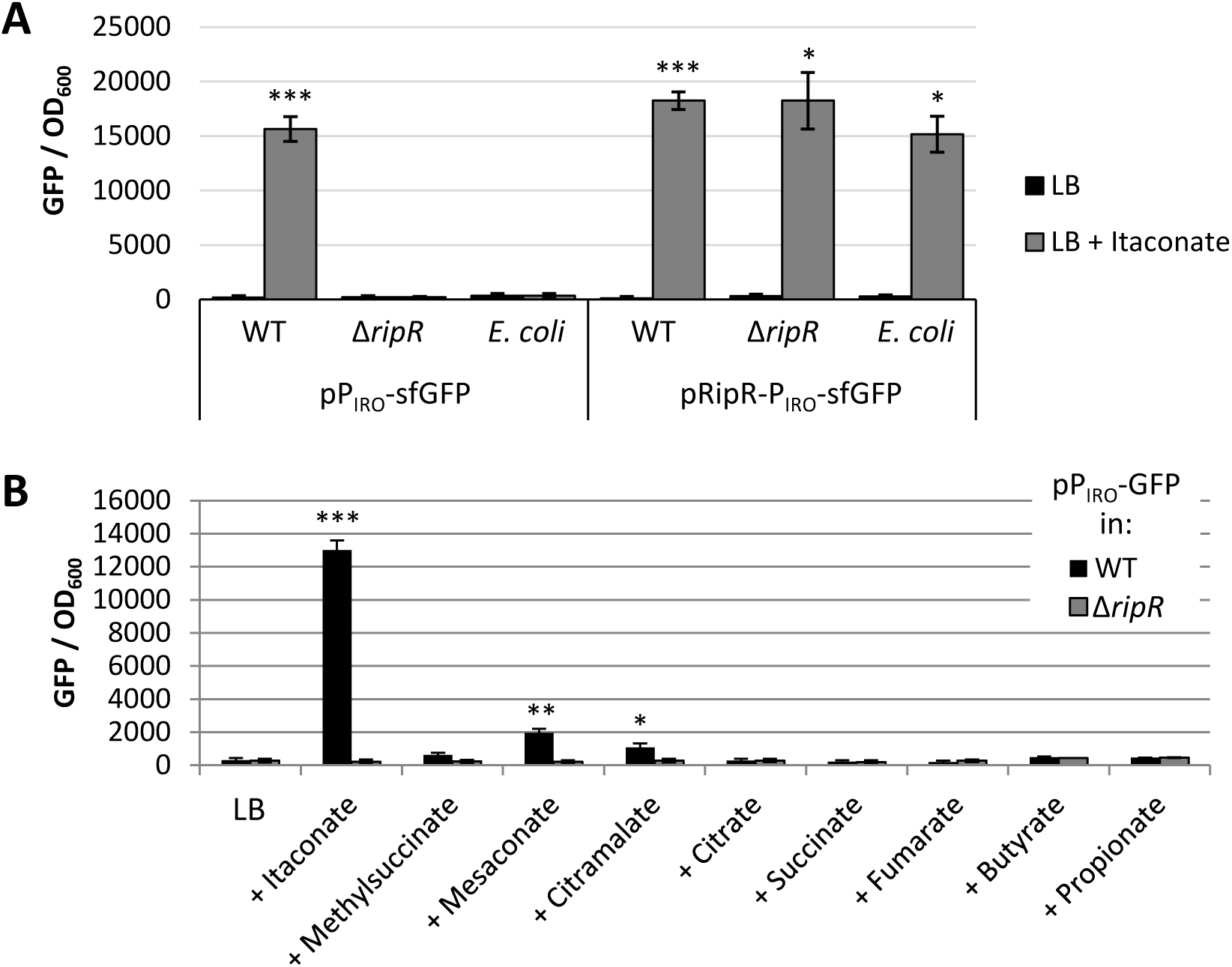
RipR is necessary and sufficient to induce P_IRO_ expression in response to itaconate. Expression of P_IRO_-sfGFP in wild-type (WT) or *ripR* knockout (Δ*ripR*) *Salmonella*, or in *E. coli*. Figures show GFP fluorescence normalized to optical density at 600nm (OD_600_) after 16h of growth in LB alone or supplemented with 0.2% itaconate (A) or similar metabolites (B). Data are the average of at least three biological replicates and error bars show one standard deviation. pP_IRO_, plasmid-borne transcriptional fusion of P_IRO_ to sfGFP; pRipR-P_IRO_, pP_IRO_ with the *ripR* gene and its native promoter included on the plasmid. A Games-Howell ANOVA was conducted comparing with and without itaconate for each strain (A), or comparing WT to Δ*ripR* (B) for each added metabolite. *, p < 0.05; **, p < 0.01; ***, p < 0.001.

To assess if the IRO promoter is induced specifically by itaconate, we examined induction of the P_IRO_ reporter plasmid in media supplemented with a panel of similar metabolites. While mesaconate, citramalate and methylsuccinate (in order of induction strength) slightly induced expression, induction by itaconate was drastically more pronounced, suggesting that it is the principal inducer (Figure 3B). Notably, similar results were obtained in complex media (LB) and in MOPS minimal media with either glucose or glycerol as a carbon source, suggesting that the induction only requires itaconate and not additional factors in the media (Figure S2). Furthermore, induction by itaconate occurred in a dose dependent manner indicating that the reporter can be used to semi-quantitatively assess itaconate concentrations encountered by the bacteria (Figure S3).

### Itaconate import is independent of the dicarboxylate transporter DctA

Another important question was whether or not itaconate required the primary aerobic dicarboxylate transporter, DctA, for import into the cell. We compared itaconate-mediated induction of the P_IRO_-sfGFP reporter in wild-type or Δ*dctA Salmonella*. Interestingly, we found that the IRO promoter was still heavily induced in the *dctA* mutant in the presence of itaconate (Figure S4). This demonstrates that the DctA dicarboxylate transporter is not required for itaconate-mediated induction of P_IRO_, suggesting that itaconate import is independent of *dctA*.

### Itaconate is bactericidal at low pH

Previous work has demonstrated that itaconate can inhibit the function of the glyoxylate shunt enzyme, AceA, and act as a bacteriostatic agent when bacteria rely on carbon sources such as acetate that require this pathway ^11–13^. It has also been suggested that itaconate can inhibit bacteria by influencing media pH and at least one publication has demonstrated that it can have bactericidal activity ^15,16^. To clarify this later phenotype we hypothesized that the dicarboxylic acid chemistry of itaconate would allow it to act in a bactericidal fashion at low pH by acting as a proton shuttle. In brief, the carboxyl groups of itaconate (pK_a_ = 5.5 and 3.8) protonate and lose their charge at lower pH allowing them to traverse the bacterial membrane and release the protons in the more neutral pH of the cytoplasm, thereby exacerbating acid stress (Figure 1A).

To emulate the intracellular conditions that *Salmonella* may encounter in a *Salmonella* containing vacuole (SCV) of a macrophage we added itaconate to LPM media and then acidified to pH 4.4, 5.0, or 5.8 to cover a range from the most acidic to more regular estimates of SCV pH ^26–28^. Indeed we found that wild-type *Salmonella* showed a 1000-fold decrease in survival after 3h hours at pH 4.4 with itaconate (Figure 1B). This lethality was alleviated at higher pH and also occurred using a similar dicarboxylic acid, succinate. Importantly, the bactericidal effect was also dependent on the presence of itaconate or succinate, as pH 4.4 LPM did not kill *Salmonella* in the absence of a dicarboxylic acid (Figure S1A). Interestingly, deletion of the entire IRO (STM3120-STM3117) or *aceA* had no effect, but deletion of the general stress response sigma factor, RpoS (σ^32^, σ^S^), exacerbated *Salmonella’s* sensitivity at both pH 4.4 and 5.0 (Figures 1C and S1B). Cumulatively, these data demonstrate that itaconate or other dicarboxylic acids can act in a bactericidal fashion under acidic conditions by exacerbating acid stress.

### The *Salmonella* IRO does not significantly contribute to short-term survival in a mouse macrophage cell line

The inhibitory effect of itaconate on AceA, its bactericidal activity under acidic conditions, and the synthesis of itaconate in macrophage combine to support the concept that these immune cells may be employing itaconate as an antibacterial compound. As a successful pathogen, *Salmonella* has adapted to survive in activated macrophage and multiple previous works have examined how IRO genes may influence *Salmonella* survival and virulence ^21–23^. In our hands, we found no significant reduction in survival of Δ*IRO* or Δ*ripR* strains in the mouse J774 macrophage cell line (Figure S5). When the macrophages were pre-stimulated with IFN-γ, there was a slight reduction in survival relative to wild-type, but this was not significant when compared to an *aceA* mutant that showed no survival defect. In contrast, the growth of a *phoP* deletion control strain was inhibited by macrophages even without IFN-γ.

### *Salmonella* encounter itaconate in the phagosomes of mouse and IFN-γ-stimulated human macrophage

Multiple high throughput studies have demonstrated that the IRO genes are induced heavily in mouse macrophage ^18–20^. Additional studies have identified itaconate in both mouse and human macrophage but the mouse cells appear to produce significantly more of the metabolite ^13,14^. Furthermore, it has been demonstrated that interferon-β (IFN-β) and IFN-γ can stimulate itaconate production in mouse macrophage ^15,29–31^.

To examine itaconate levels encountered by intracellular bacteria inside macrophage, we employed our P_IRO_-sfGFP reporter plasmid as a biosensor. By including a constitutively expressed mCherry gene on the same plasmid, we could microscopically observe individual bacteria inside macrophage and obtain semi-quantitative data by generating a GFP/mCherry fluorescence ratio as an indicator of P_IRO_ induction and accordingly itaconate levels. Using this system, we observed strong induction of the P_IRO_ promoter for wild-type *Salmonella* in J774 mouse macrophage and this signal was absent in the Δ*ripR* control (Figure 4). Furthermore, the response could also be observed in *E. coli* if RipR was encoded on the same plasmid, demonstrating that bacteria that are poorly adapted to intracellular survival also encounter itaconate in macrophage.

**Figure 4:**
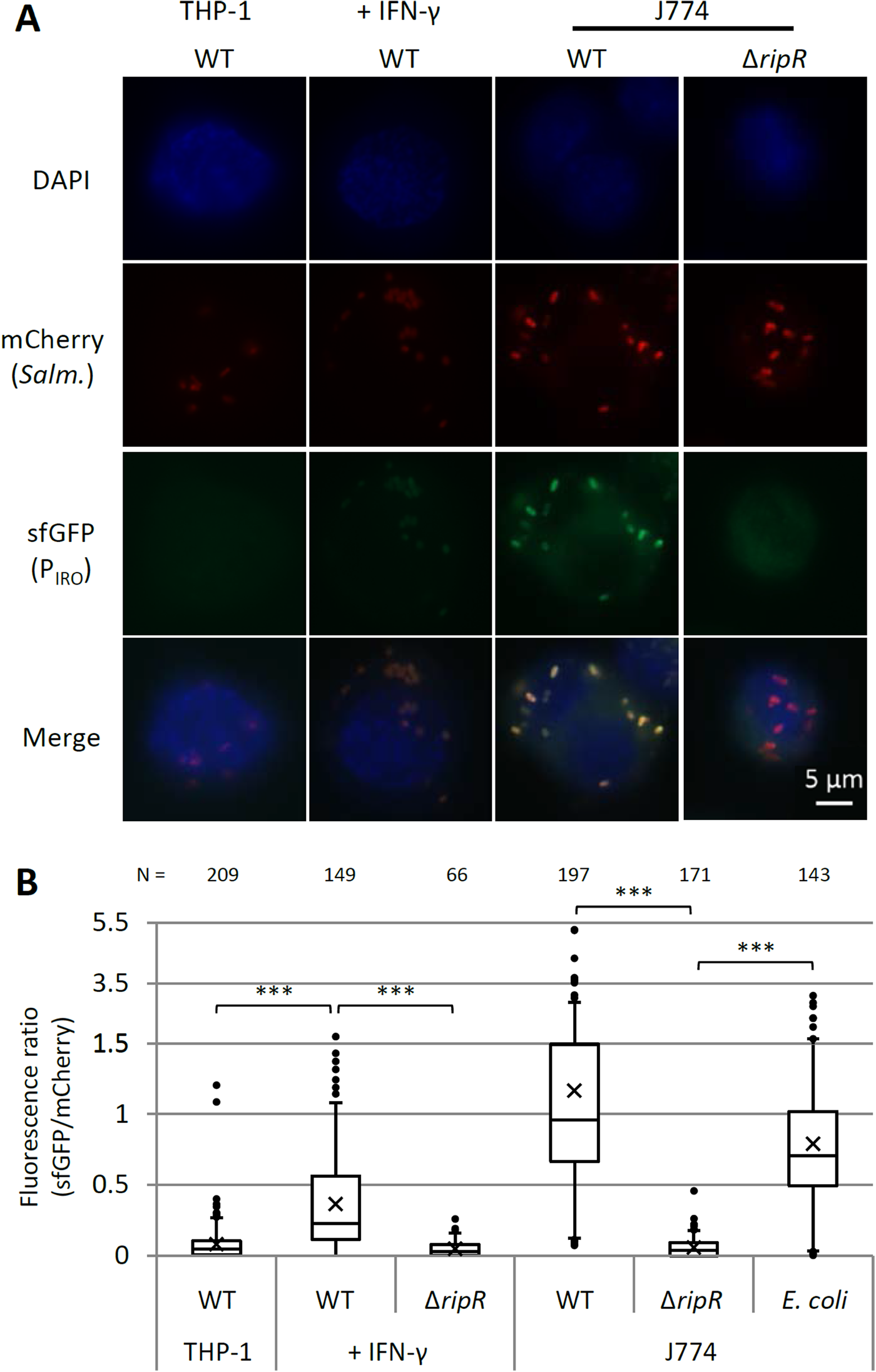
P_IRO_ is activated in J774 macrophage and THP-1 macrophage stimulated with IFN-γ. Expression of P_IRO_-sfGFP in intracellular wild-type (WT) or *ripR* knockout (Δ*ripR*) *Salmonella* containing the pICM-P_IRO_ plasmid (constitutive mCherry expression). For expression in *E. coli*, the *ripR* gene was also encoded on the reporter plasmid. ‘+ IFN-γ’ samples were pre-treated with human IFN-γ. **A)** Representative fluorescence microscopy images. **B)** Relative fluorescence quantification of individual bacterial particles at 8h post-infection. Number of bacteria quantified is indicated above and totalled from at least two biological replicates (≥ 3 for all WT *Salmonella* samples). Boxes indicate first and third quartiles; central line, median; X, mean; whiskers, 95^th^ percentile; dots, all non-zero data points outside 95^th^ percentile. The y-axis changes scale at 1.5 to better show outliers. A Games-Howell ANOVA was conducted comparing indicated samples. ***, p < 0.001.

In contrast to the mouse cell line, unstimulated human THP-1 monocytes showed negligible itaconate levels as very few of the bacterial reporters showed any green fluorescence above background levels (Figure 4). The bacteria did express the constitutive mCherry and could be observed in the macrophage, suggesting that the lack of green fluorescence was not due to decreased bacterial survival or protein expression. Stimulation of THP-1 cells with IFN-γ (M1 activation), however, led to a significant increase in the green fluorescence of the reporter bacteria in a RipR-dependent fashion. In contrast, itaconate levels in THP-1 cells induced with IL-4 and IL-13 (M2 activation) resembled unstimulated cells (Figure S6).

## Discussion

In this work we demonstrate that itaconate becomes bactericidal at acidic pH, suggesting an additional mechanism for itaconate to act as an antibacterial metabolite beyond inhibition of AceA. Thus, elevated itaconate levels in macrophage may act to inhibit bacterial metabolism while also exacerbating acid stress on microbes in the phagosome. Protonation of itaconate under acidic conditions may also grant it increased access to the bacterial cytoplasm where de-protonation would trap the charged form close to its AceA target. This organic acid killing effect has been demonstrated previously, including in a recent work showing propionate inhibition of *Salmonella* in mice ^32,33^. Of note, we also find that bacterial killing occurs with succinate, a metabolite similar to itaconate that similarly increases in concentration in activated macrophage ^34^. Our findings that these dicarboxylic acids can kill *Salmonella* at pH 4.4 but not higher may contribute to why *Salmonella* manipulates the SCV to maintain a pH closer to 5.0 in order to avoid this organic acid stress. Moreover, they imply that organic acids such as itaconate and succinate may contribute to the antibacterial activity of acidified phagosomes.

The antibacterial potential of itaconate, its synthesis in activated macrophage, and the localization of IRG1 to bacteria-containing vacuoles, support its potential role as a weapon against intracellular bacteria. Here we examined survival of *Salmonella* IRO or *ripR* deletion strains in the mouse macrophage J774 cell line but saw no significant decrease in survival, similar to a recent study examining SPI-13 in RAW264.7I macrophage ^24^. However, in that work, Espinoza *et al*. discovered that SPI-13 does play a role in *Salmonella* internalization into mouse – but not human – macrophage ^24^. Combined with previous works showing reduced survival of IRO mutants in mice, this operon may play a more significant survival role in the context of infection in animals that produce copious amounts of itaconate ^20–23^.

We find *Salmonella* responds to itaconate *in vitro* and intracellularly by strongly inducing an operon encoding itaconate degradation proteins. This response is largely specific to itaconate and is entirely dependent on the neighbouring gene (*STM3121* in the prototypical *Salmonella enterica* Sv. Typhimurium strain LT2), which we propose to rename *ripR* (‘rip operon regulator’) to conform to earlier nomenclature. The *ripR* gene product is predicated to be a LysR family transcriptional regulator, suggesting that itaconate induces IRO expression by interacting directly with the substrate binding domain of RipR to activate it. Moreover, we find that RipR is sufficient for itaconate induction of the P_IRO_ promoter in *E. coli*, demonstrating its potential as a novel inducible expression system with over 50-fold higher transcription in the presence of the inexpensive and readily available inducer. A limitation of this expression system would be a requirement for growth on carbon sources independent of the glyoxylate shunt and also growth at neutral or alkaline pH, as we demonstrate that itaconate is bactericidal under acidic conditions. However, for many studies these conditions are met, adding P_IRO_ to the repertoire of available inducible promoter systems.

Using our P_IRO_-sfGFP itaconate biosensor, we showed a pronounced response in unstimulated mouse macrophage whereas no induction was observed in the THP-1 human macrophage cell line without stimulation, suggesting that these cells are not producing itaconate to the same degree. Interferon has previously been demonstrated to stimulate itaconate production in mouse macrophage and we found that our biosensor was induced in human cells stimulated with IFN-γ ^15,29–31^. Thus, while the human cell line was able to produce itaconate, it required auxiliary induction to do so and still produced less than the uninduced mouse macrophage. While it is possible that this reflects an artifact of the cell lines employed, it aligns well with previous works that quantified itaconate in both mouse and human cells ^13,14^. Furthermore, Espinoza *et al*. recently determined that SPI-13 is abundant in generalist *Salmonella* serovars but not human-restricted ones (which instead encode SPI-8), suggesting that low itaconate levels in humans render the IRO dispensible in these strains ^24^.

A recent study demonstrated that small molecules can inhibit the activity of the IRO proteins and sensitize *Salmonella* to itaconate inhibition in minimal media ^35^. Such drugs could also sensitise other bacteria to itaconate including *Yersinia, Pseudomonas* and *Mycobacteria* species, which also encode an IRO ^17,35^. Moreover, if human-restricted pathogens lack an IRO because human cells truly produce less itaconate, then they are potentially sensitive to it and itaconate itself could potentially be used as an antimicrobial against them. Our biosensor could prove invaluable in such studies for determining how much itaconate the bacteria are encountering, and the self-sufficiency of the biosensor allows it to be employed in a variety of species, providing added versatility.

In summary, here we present data that itaconate can act as a bactericidal metabolite at acidic but physiologically relevant pH. We identify the regulatory mechanism of an itaconate response operon in *Salmonella* and employ its promoter as a novel biosensor of relative itaconate concentrations in macrophage phagosomes. Finally, we provide further evidence that IFN-γ stimulates itaconate synthesis and moreover that human cells produce less of the metabolite than their mouse equivalents.

## Materials and Methods

### Bacterial strains and plasmids

All *Salmonella* strains used in experiments are derivatives of *Salmonella enterica subsp. enterica* serovar Typhimurium (*S*. Typhimurium) strain 14028s. As described previously, lambda red recombination and subsequent P22 phage transduction was used to generate all of the gene knockout mutants in this background ^36–38^. To allow for subsequent recombinations, the antibiotic resistance cassette was removed from the chromosome using the pCP20 plasmid encoding FLP recombinase ^39^. The heat-unstable pCP20 plasmid was eliminated by passaging overnight at 42°C and loss was confirmed by antibiotic treatment.

A reporter fusion of the IRO promoter to sfGFP (P_IRO_-sfGFP) was generated using Gibson cloning to insert the 333bp upstream of the STM3120 start codon (thereby including 25bp upstream of the predicted -35 box and the 5’ untranslated region) into the pXG10sf plasmid (replacing the existing promoter)^40–42^. For the reporter construct including *ripR*, the same region was extended to 1570bp upstream of the STM3120 start codon to include the entire STM3121 ORF and a predicted transcriptional terminator following it. For fluorescence microscopy, constitutively expressed (PLtet0-1 promoter) mCherry was inserted into a transcriptionally independent region of the same plasmid. This variation of the plasmid was renamed ‘independent constitutive mCherry’ or pICM.

### Metabolite induction of P_IRO_ assay

Induction of the *Salmonella* P_IRO_ promoter was assessed using a transcriptional fusion to sfGFP in either the pXG10sf or pICM plasmids. Data from the two plasmids were combined as the inducible region is identical and the plasmids only differ in the constitutively active mCherry expressed independently on pICM. Overnight LB cultures were used to inoculate (1/200 dilution) either LB or MOPS minimal media containing 0.2% of the indicated carbon source. Itaconate or other metabolites at neutral pH were supplemented to a concentration of 0.2%. Of note, for salts and hydrates the final 0.2% concentration reflects the percent of the carbon source itself; e.g. 0.2% succinate was made as 0.47% sodium succinate (dibasic) hexahydrate. Growth was conducted in a TECAN Infinite M200 plate reader at 37°C with shaking and OD_600_ and GFP fluorescence (475nm and 511nm excitation and emission wavelengths respectively) were read every 15 minutes. For clarity, bar graphs show fluorescence at 16h post inoculation. Chloramphenicol was included in all media at a concentration of 20μg/ml to maintain the plasmids.

### Acidified media survival

LPM media was made as described previously ^43,44^. Succinate or itaconate were added to 0.4% and the pH was then adjusted to 4.4, 5.0 or 5.8 as indicated. LB overnight cultures were resuspended in acidified media to an OD of 0.1 and incubated in a 37°C water bath. At time points, samples were taken, serial diluted and plated for colony forming units (CFU).

### Intra-macrophage survival

The THP-1 human monocyte cell line and the J774 mouse macrophage cell line were maintained in RPMI Medium 1640 (with L-glutamine) supplemented with 10% FBS and 1% Glutamax and grown at 37°C and 5% CO_2_. THP-1 cells were seeded in 96-well plates at 50,000 per well with 50nM PMA (phorbol 12-myristate 13-acetate) added to induce differentiation to adherent macrophage. After 48h, media was replaced with no-PMA growth media overnight with 100 U/ml human IFN-γ or IL-4 and IL-13 added if indicated. For J774 macrophage the cells were seeded in 96-well plates at 50,000 per well overnight with 100 U/ml mouse IFN-γ added if indicated. *Salmonella* in RPMI were added onto seeded cells at a multiplicity of infection (MOI) of approximately 20:1 and centrifuged for 10 minutes at 1000 rpm to maximize cell contact. After centrifuging the samples were incubated at 37°C and 5% CO_2_ (time 0). After 30 minutes, cells were washed three times with PBS followed by fresh media containing 100 μg/ml gentamicin to kill extracellular *Salmonella*. At 2 hours the media was replaced with media containing gentamicin at 10 μg/ml. At timepoints, intracellular bacteria were recovered using PBS containing 1% Triton X-100 and vigorous pipetting. Samples were serially diluted and five 10μl spots were plated for CFU counting. Each sample included three separate wells as technical replicates (a total of 15 × 10μl spots counted per biological replicate).

### Fluorescence microscopy

Fluorescence microscopy was conducted similarly to the macrophage survival assay with some exceptions: Cells were seeded in 24-well plates containing glass coverslips at 125,000 per well. Bacteria were infected at an MOI of approximately 100 to maximize the instances of macrophage containing bacteria. At timepoints the media was removed and cells were washed three times with PBS. They were then fixed for 10 minutes at room temperature in PBS + 4% paraformaldehyde (PFA). Following three more PBS washes the cells were permeabilized for 10 minutes in PBS + 0.2% Triton X-100 + 1% BSA. Coverslips were washed again, mounted on slides using 3μl mounting media containing DAPI and allowed to dry overnight in the dark. Slides where viewed using a Zeiss Observer.z1 microscope using a 100x oil immersion objective and the Zeiss Zen microscopy software. Images were taken with a Zeiss Axiocam 506 mono camera mounted on the microscope. For all samples a 2s exposure was used for mCherry and 1s exposure for sfGFP. ImageJ software was employed for quantification to calculate fluorescence intensities in the red and green channels relative to a neighbouring background region for each bacterium and a GFP/mCherry ratio was generated.

## Acknowledgements

We would sincerely like to thank Dr. Scott Gray-Owen and members of his lab, in particular Dr. Ryan Gaudet, for their generous donation of technical expertise, macrophage cells lines, and use of their equipment. WWN was supported by an Operating Grant from the Canada Institutes for Health Research (MOP-86683) and a Natural Sciences and Engineering Research Council (NSERC) of Canada Grant (RGPIN 386286-10). SJH was supported by an NSERC Vanier Canada Graduate Scholarship.

## Supplemental Figures

**Figure S1:**
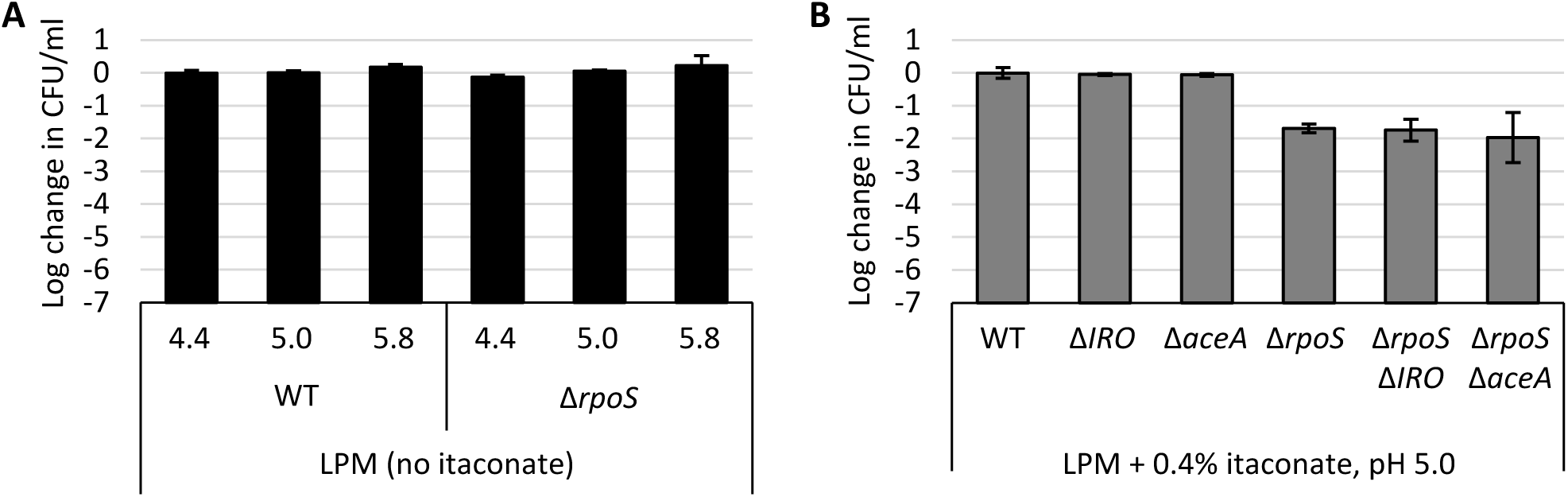
pH 4.4 without itaconate was not bactericidal. **A)** Survival (relative to 0h time point) of wild-type (WT) or *rpoS* mutant *Salmonella* after 3h in LPM media adjusted to pH 4.4, 5.0, or 5.8 as indicated. **B)** Survival of *Salmonella aceA* and itaconate response operon (IRO) mutants in LPM + itaconate media at pH 5.0. All data are the average of at least three biological replicates and error bars show one standard deviation.

**Figure S2:**
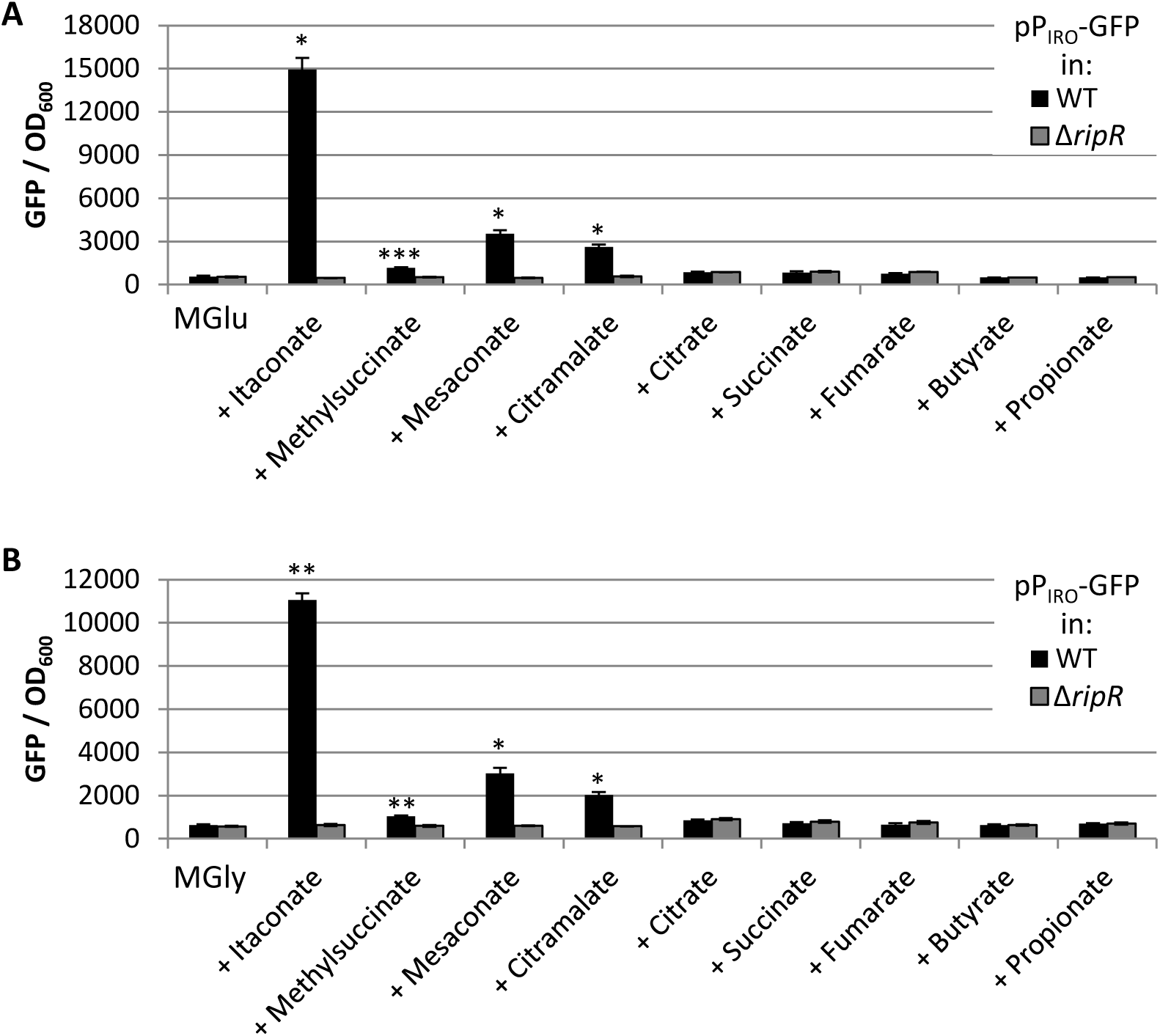
Itaconate induction of P_IRO_ occurs in minimal media. Expression of P_IRO_-sfGFP in wild-type (WT) or *ripR* knockout (Δ*ripR*) *Salmonella* as in Figure 1 except grown in MOPS minimal media with 0.2% Glucose (**A**; MGlu) or Glycerol (**B**; MGly) instead of LB. Data show GFP fluorescence normalized to OD_600_ after 16h of growth and are the average of at least three biological replicates. Error bars show one standard deviation. pP_IRO_, plasmid-borne transcriptional fusion of P_IRO_ to sfGFP. A Games-Howell ANOVA was conducted comparing WT to Δ*ripR* for each added metabolite. *, p < 0.05; **, p < 0.01; ***, p < 0.001.

**Figure S3:**
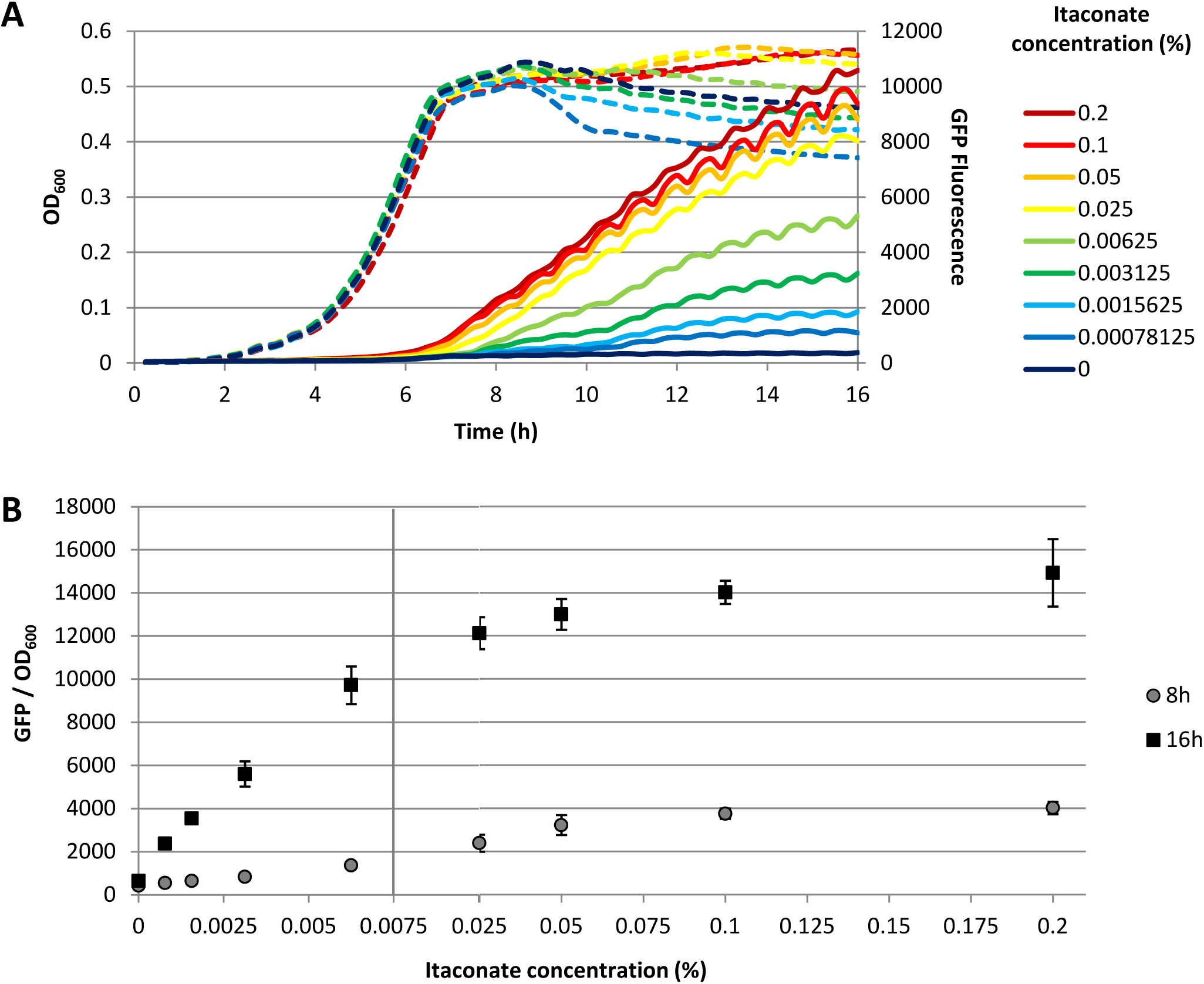
P_IRO_ is induced by itaconate in a dose-dependent manner. **A)** Example replicate showing *Salmonella* optical density (dashed lines) and P_IRO_-sfGFP fluorescence (solid lines) for a range of itaconate concentrations across 16h of growth. All conditions were in MOPS minimal media with 0.2% glucose as a carbon source and indicated concentration of itaconate (%, g/100ml). **B)** Average P_IRO_-sfGFP fluorescence (normalized to OD_600_) at 8h or 16h across two biological replicates. Error bars indicate one standard deviation. The x-axis changes scale at 0.0075% to show lower concentrations more clearly.

**Figure S4:**
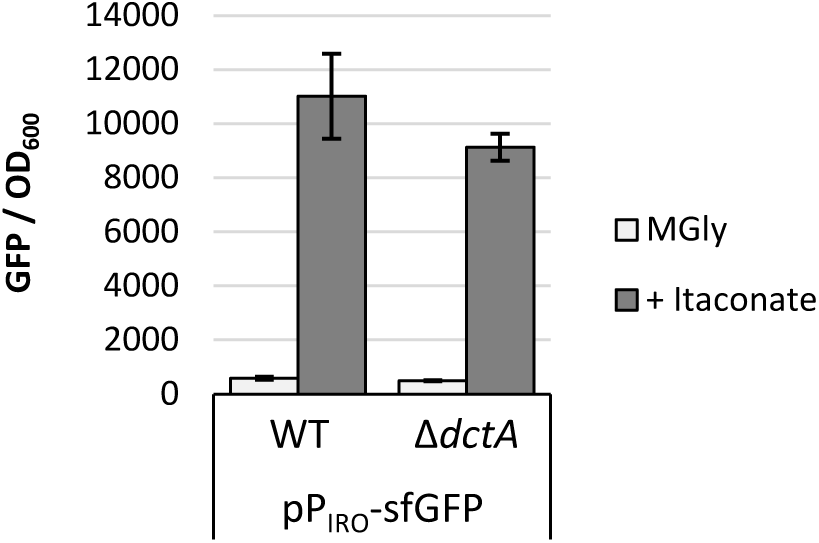
DctA is not required for the response to itaconate. Expression of P_IRO_-sfGFP in wild-type (WT) or *dctA* knockout (Δ*dctA*) *Salmonella*. Figure shows GFP fluorescence normalized to optical density at 600nm (OD_600_) after 16h of growth in MOPS minimal media with 0.2% glycerol as the carbon source. Data are the average of at least two biological replicates and error bars show one standard deviation. pP_IRO_-sfGFP, plasmid-borne transcriptional fusion of P_IRO_ to sfGFP.

**Figure S5:**
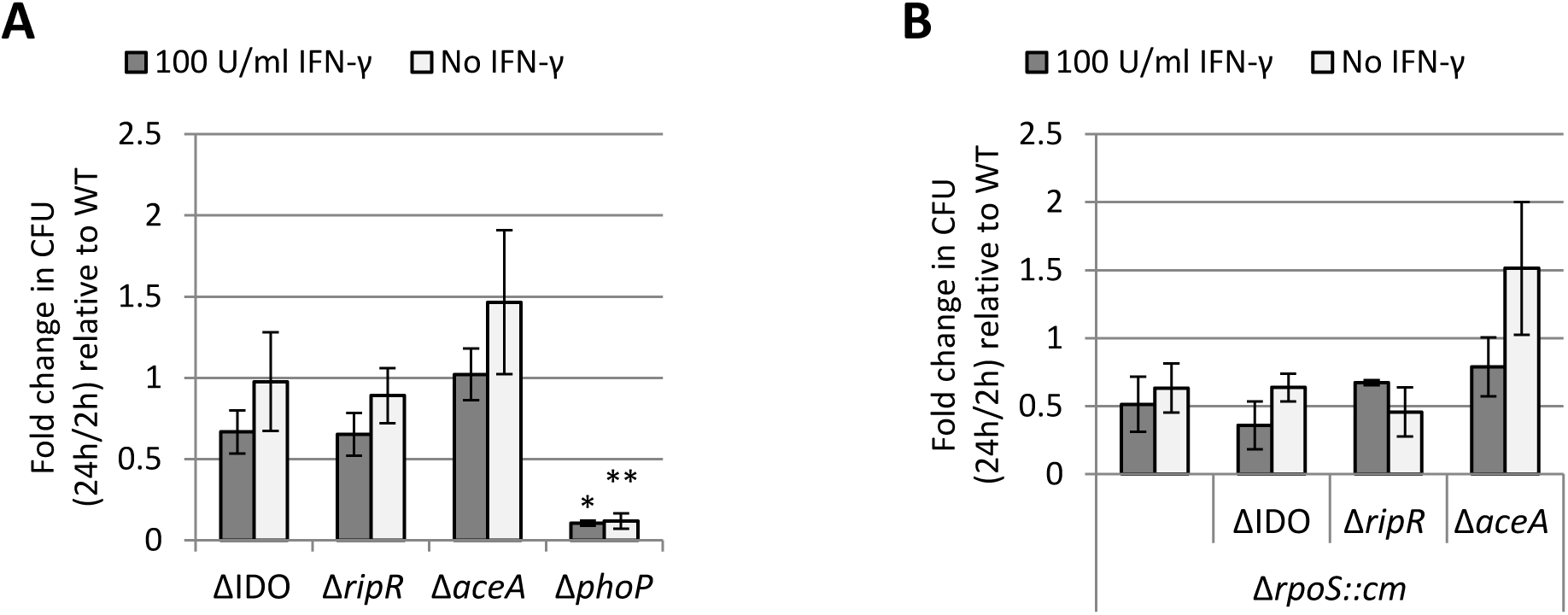
Survival of *Salmonella* mutants in J774 macrophage with and without IFN-γ stimulation. relative to wild-type. The fold change in CFU recovered (24h relative to 2h post-infection) is shown and data from each replicate was normalized to the 24h/2h ratio obtained for wild-type *Salmonella*. Columns show the average of three biological replicates and error bars show one standard error of the mean. Since samples were normalized to wild-type in each replicate, a one-way ANOVA with Sidak’s multiple comparison test was conducted comparing each strain to the Δ*aceA* strain (A) or to the Δ*rpoS* strain (B). Only the Δ*phoP* mutant was found to have a p < 0.05. *, p<0.05; **, p < 0.01.

**Figure S6:**
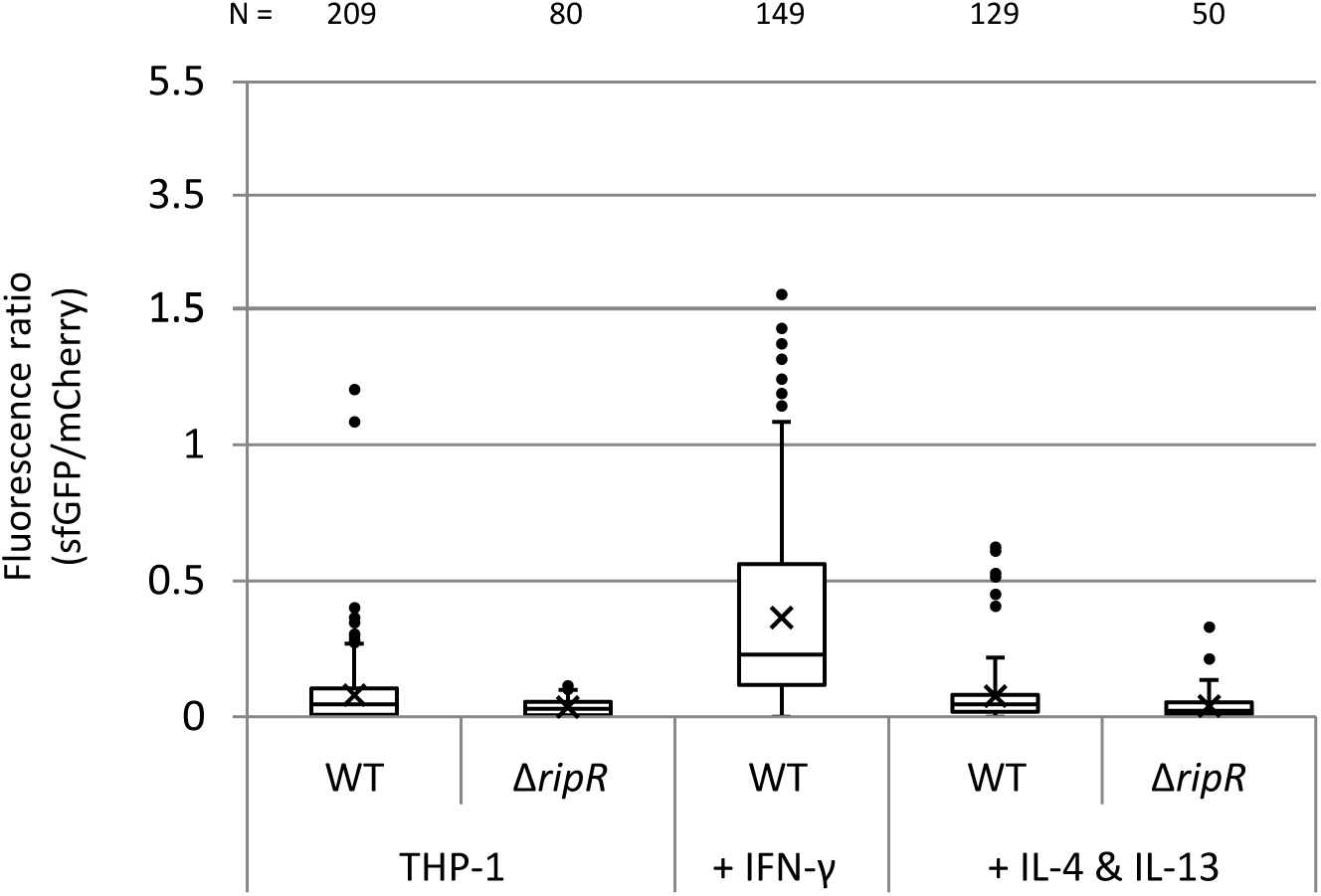
P_IRO_ shows little-to-no activation in THP-1 macrophage stimulated with IL-4 and IL-13. As in Figure 4, expression of P_IRO_-sfGFP in intracellular wild-type (WT) or *ripR* knockout (Δ*ripR*) *Salmonella* containing the pICM-P_IRO_ plasmid (constitutive mCherry expression). THP-1 macrophage were pre-treated with no stimulant, human IFN-γ, or IL-4 and IL-13 and infected with *Salmonella* for 8h. Data for WT *Salmonella* in THP-1 with no treatment or IFN-γ are shown in Figure 4 and are shown again here for comparison. As in Figure 4, data shows relative fluorescence quantification of individual bacterial particles. Number of bacteria quantified is indicated above and totalled from at least three biological replicates. Boxes indicate first and third quartiles; central line, median; X, mean; whiskers, 95^th^ percentile; dots, all non-zero data points outside 95^th^ percentile. The y-axis changes scale at 1.5 to better show outliers.

